# Morphological peculiarities of DNA-protein complexes in dormant *Escherichia coli* cells, subjected to prolonged starvation Condensation of DNA in dormant cells of *Escherichia coli*

**DOI:** 10.1101/2020.03.27.011494

**Authors:** Natalia Loiko, Yana Danilova, Andrey Moiseenko, Vladislav Kovalenko, Ksenia Tereshkina, Galina I. El-Registan, Olga S. Sokolova, Yurii F. Krupyanskii

**Affiliations:** Department of Structure of Matter, Semenov Federal Research Center of Chemical Physics, RAS, Moscow, Russia; Department of Microbiology, Federal Research Center ‘Fundamentals of Biotechnology’ RAS, Moscow, Russia; Department of Biology, Lomonosov Moscow State University, Faculty of Biology, Moscow, Russia

**Author notes:** Corresponding authors: (Yu.F.K), (O.S.S).

## Abstract

One of the adaptive strategies for the constantly changing conditions of the environment utilized in bacterial cells involves the condensation of DNA in complex with the DNA-binding protein, Dps. With the use of electron microscopy and electron tomography, we observed several morphologically different types of DNA condensation in dormant *Escherichia coli* cells, namely: *nanocrystalline, liquid crystalline*, and the *folded nucleosome-like*. We confirmed the presence of both Dps and DNA in all of the ordered structures using EDX analysis. The comparison of EDX spectra obtained for the three different ordered structures revealed that in *nanocrystalline* formation the majority of Dps protein is tightly bound to nucleoid DNA. We demonstrated that the population of the dormant cell is structurally heterogeneous, which allows cells to respond flexibly to environmental changes. It increases the ability of the whole bacterial population to survive under extreme stress conditions.

## Introduction

Living organisms survive in constantly changing environmental conditions, due to the universal strategies of adaptation to various stresses based on structural, biochemical, and genetic rearrangements. The universal adaptive response of microorganisms to starvation is of specific interest, because it often coincides with the antibiotic’s resistance of pathogenic bacteria; this represents one of the most important medical problems in the world to date [1].

Adaptive molecular strategies ensure the ability of microorganisms to survive in environments significantly different from those, optimal for their growth, and have been the subject of active research over the past years [2–8]. Such adaptive strategy often launch the increasing synthesis (up to 150-200 thousand copies) of a ferritin-like protein Dps (<u>D</u>NA-binding <u>p</u>rotein of <u>s</u>tarved cells) [9–12]. Dps carries on a regulatory and protective role within *Escherichia Coli* (*E. coli)* cells [13]. Its structure [14] and interactions with DNA were recently excessively studied *in vitro* [15,16], and *in silico* [17,18]. Increased synthesis of Dps in *E. coli* cells occurs in the stationary phase, under starving conditions, allowing for the protection of DNA from oxidative stress, heat, acid, alkaline shock, toxic effects of heavy metals, antibiotics, UV radiation, etc. [9,15,19–21]. The protective functions of Dps are carried out through condensation of DNA into “*biocrystalline*” or “*in cellulo nanocrystalline*” structures [2–7,19,22-24]

It is considered that the bacterial nucleoid represents an intermediate engineering solution between the protein-free DNA packaging in viruses and protein-defined DNA packaging in eukaryotes [25]. In diluted solutions, the diameter of a DNA double-helix is about 2 nm, while its length may reach up to several centimeters. In a normal bacterial cell, the circular DNA is located within a well-defined region, called the nucleoid, which fills only 15% to 25% of the total volume. Under physiological conditions, DNA condensation leads to a dramatic decrease in the volume occupied by the DNA in the cytoplasm [26]. Polymer physicists often call this process a “coil-globule” transition [27]. By using 3C- and Hi-C-based techniques, it has been demonstrated that DNA spatial organization is different in each cell [28,29]. This corresponds to the heterogeneity of cells within a population. The heterogeneity of cells allows for a flexible response to environmental changes and helps to survive in stressful situations [5].

Considering everything priory mentioned, we may expect to observe a somewhat different structural response to stress within different bacterial cells. For this reason, this study was devoted to the observation of morphological differences in structures, formed by condensed DNA in dormant *E. coli* cells and to the experimental investigation of the variety of DNA packaging in cells of different strains and growing conditions, under the stress of prolonged starvation.

## Materials and methods

### Bacterial strains

All bacterial strains were kindly provided by Prof. Vassili N. Lasarev, Head of the Laboratory of Gene Engineering, Federal Research and Clinical Center of Physical-Chemical Medicine, Federal Medical Biological Agency, Moscow, Russia.

1. gram-negative bacteria *E. coli* Top10;
2. Dps over-producer strain *E. coli T*op10/pBAD-DPS;
3. Dps over-producer strain *E. coli* BL21-Gold(DE3)/pET-DPS.

### Reagents used

All reagents were purchased from Sigma-Aldrich (St Louis, MO) or VWR (Solon, Ohio, USA), unless stated otherwise. LB medium (LB Broth, Miller (Luria-Bertani)) was purchased in VWR (Life Science, Lot: 18G3056107). Agar powder was obtained from Alfa Aesar GmbH and Co (Karlsruhe, Germany, Lot 10143436). Modified M9 medium contains the following (g/L): Na_2_HPO_4_ – 6; KH_2_PO_4_ – 3; NaCl – 0.5; NH_4_Cl – 0.2; MnSO_4_ – 0.0004; MgSO_4_ – 0.0025; CaCl_2_ – 0.0002; glucose – 10; pH =7.0.

### Preparation of genetically modified Strains of *E. coli* Top10/pBAD-DPS and *E. coli* BL21-Gold(DE3)/pET-DPS

The DNA fragment encoding Dps was obtained by PCR amplification of the DNA from *E. coli* K12 MG1655, using oligonucleotides dps-nde and dps-hind (http://www.uniprot.org/uniprot/P0ABT2): dps-nde 5’-GATATGAACATATGAGTACCGCTAAATTAG; dps-hind 5’-TATAAGCTTATTCGATGTTAGACTCGATAAAC.

Expression plasmids were obtained using two vectors with different transcription promoters and two recipient strains, *E. coli* pET-min and *E. coli* pBAD/Myc-His A.

The DNA fragment was introduced into the pETmin plasmid at restriction sites: NdeI and HindIII. This resulted in the production of a pET-DPS plasmid containing the DNA region encoding Dps under control of the T7 promoter.

The structure of the pBAD/Myc-His A recipient vector prevented introduction of the coding DNA fragment using restriction sites so that the recombinant protein would contain no additional amino acid sequences compared to natural Dps. The promoter was, therefore, deleted by cleavage at the BamHI and HindIII restriction sites. The excised region of the plasmid was then amplified on the pBAD/Myc-His A template, using the PB-F and PB-dpsR oligonucleotides. The Dps-encoding DNA fragment was obtained by PCR amplification of the DNA from *E. coli* K12 MG1655, using the PB-dpsF and dps-hind oligonucleotides. DNA fragments were purified by preparative electrophoresis in agarose gel, pooled, and amplified using the PB-F and PB-dpsR oligonucleotides. The DNA fragment was then introduced into the pBAD/Myc-His A plasmid, using the BamHI and HindIII restriction sites. The resultant pBAD-DPS plasmid contained the Dps-encoding DNA region under the control of an *E. coli* arabinose operon promoter.

### Dps overproduction in genetically constructed *E. coli* cells

*E. coli* strain BL21-Gold was transformed by the pET-DPS plasmid. *E. coli* strain Top10 was transformed by the pBAD-DPS plasmid. For protein expression, the medium (LB, Amp 150 mg/L, 10 mM lactose for BL21-Gold or 6.7 mM arabinose for Top10) was inoculated with a single bacterial colony and incubated with shaking for 16–18 hrs at 37°C. The cells were collected by centrifugation (5000 g, 15 min), resuspended in water, and homogenized with ultrasound. The lysate was centrifuged for 10 min at 13000 g. After the removal of the supernatant (water-soluble fraction), the pellet was resuspended in 1% SDS, heated for 5 min at 95°C, and centrifuged. The supernatant (insoluble fraction) was collected. Protein composition of the supernatants was analyzed by SDS-PAAG, according to Laemmli [24]. The total yield of recombinant Dps were assessed as high, constituting ∼50% of the total protein in the cells of producer strains. The density of the protein bands was estimated using the gel electrophoresis image analysis software GelAnalyzer.

### Preparation of dormant *E. coli* cells

#### Top10 strain

Bacteria were grown for 24 hrs at 28°C under shaking (140 rpm), in 250-mL flasks with 50 mL of LB medium. Cells were stored at 21°C, for 7 months.

#### Top10/pBAD-DPS strain

Bacteria were grown for 24 hrs at 28°C under shaking (140 rpm), in 250-mL flasks with 50 mL of either LB or modified M9 medium, supplemented with 150 μg/mL ampicillin. Dps overproduction was induced by1 g/l of arabinose in the linear growth phase. After that, cells were stored at 21°C for 7 months.

#### BL21-Gold(DE3)/pET-DPS

Bacteria were grown for 24 hrs at 28°C under shaking (140 rpm), in 250-mL flasks with 50 mL of modified M9 medium, supplemented with 150 μg/mL ampicillin with decreased ammonium nitrogen content. Dps overproduction was induced by 10 mM of lactose in the linear growth phase. The cells were stored for 7 months at 21°C.

**Light microscopy** was carried out using a Zetopan light microscope (Reichert, Austria), under phase contrast.

**The number of vegetative and dormant cells** was determined from the colony forming unit (CFU). Number obtained by plating diluted cell suspensions on agar containing (2%) LB media.

### Sample preparation for electron microscopy (EM)

Cells were fixed with 2% glutaraldehyde for 5 hrs and postfixed with 0.5 % paraformaldehyde; washed with a 0.1 M cacodylate buffer (pH=7.4), contrasted with a 1% OsO_4_ solution in a cacodylate buffer (pH=7.4), dehydrated in an increasing series of ethanol solutions, followed by dehydration with acetone, impregnated, and embedded in Epon-812 (in accordance with manufacturer’s instructions). Ultrathin sections (100 – 200 nm thick) were cut with a diamond knife (Diatome) on an ultramicrotome Ultracut-UCT (Leica Microsystems), transferred to copper 200 mesh grids, covered with formvar (SPI, USA), and contrasted with lead citrate, according to the Reynolds established procedure [31]. For analytical electron microscopy study, contrasting was in some cases omitted.

### Transmission electron microscopy (TEM)

Ultra-thin sections were examined in transmission electron microscopes JEM1011 and JEM-2100 (Jeol, Japan) with accelerating voltages of 80 kV and 200kV, respectively, and magnification of x13000-21000. Images were recorded with Ultrascan 1000XP and ES500W CCD cameras (Gatan, USA). Tomograms were obtained from semi-thick (300-400 nm) sections using the *Jeol Tomography* software (Jeol, Japan). The tilting angle of the goniometer ranges from −60° to +60° (with a permanent step of 1 degree). A series of images were aligned by the Gatan Digital Micrograph (Gatan, USA) and then recovered with the back-projection algorithm in IMOD4.9. 3D sub-tomograms were visualized in the UCSF Chimera package [32].

**Analytical electron microscopy** was carried out on an analytical transmission electron microscope JEM-2100 (Jeol, Japan), equipped with a bright field detector for scanning transmission electron microscopy (SPEM) (Jeol, Japan), a High Angular Angle Dark Field detector (HAADF) (Gatan, USA), an X-Max 80 mm^2^ Silicon Drift Detector (Oxford Instruments, Great Britain), and a GIF Quantum ER energy filter (Gatan, USA). Scanning transmission EM (STEM) and TEM modes were used. The STEM probe size was 15 nm. Energy-dispersive X-ray (EDX) spectra collection and element analyses were performed in the INCA program (Oxford Instruments, Great Britain).

### Statistical analysis

To obtain statistically reliable data and to prove the reproducibility of the experiment data, each cell line was cultivated in three independent experiment series. The type of condensed DNA structure was determined visually, after analyzing at least 20 informative micrographs of each sample, that contained around 8 – 25 viable cells. Thus, the number of cells analyzed ranged from 450 to 1500 (in all of the experiments combined). Cells with a condensed DNA structure, that could not be reliably characterized, were not considered.

## Results and discussion

Dormant cells are formed by bacteria depleted from nutrients for long periods of time [4,5,24]. Most cells (up to 99.98%) in long-term starving populations undergo autolysis. The remaining cells develop into dormant forms, which differ significantly in structural organization from vegetative cells. At stress conditions, additional condensation of DNA (comparing to the physiological conditions) proceeds. The mechanism of condensation of DNA in dormant bacterial cells is still obscure.

### Dps levels increase in cells that survived starvation and stress development

The levels of Dps expression in bacterial cells after 6, 24, 48 hrs, 2 and 7 months of cultivation have been tested using the SDS-PAGE in 10% PAAG (Fig. 1A). Experiments have shown that the biomass averaged specific levels of Dps at 48 hrs of starvation increased more than twice, comparing to protein levels in stationary cells (24 hrs), and continued to grow for up to 7 months of starvation (Fig. 1B).

**Figure 1.**
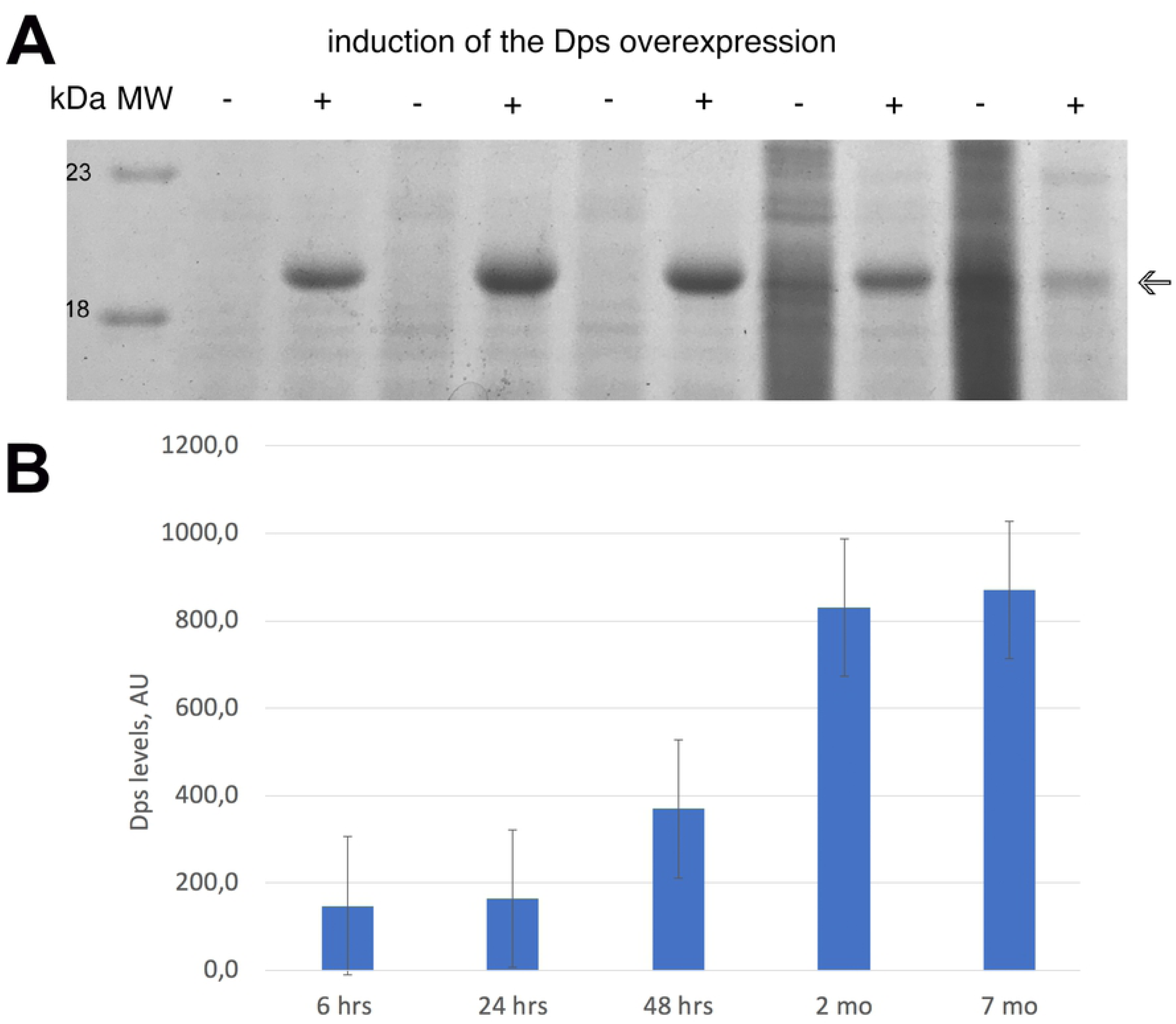
Dps content in *E. coli* cells, upon prolonged starvation. (A) SDS-PAAG electrophoresis of the biomass extract. The induction of Dps overproduction was performed at the beginning of linear growth (at 3.5 hr). Arrow – Dps; (B) The averaged amount of Dps (normalized per cell number) in *E. coli* Top10/pBAD-DPS population of cells at different intervals during starvation.

This suggests that protein synthesis in cells continues even after the termination of active growth during the transition to the stationary stage, which has been demonstrated previously [33]. The heterogeneity of the bacterial population, at this stage, led to differences in the amount of Dps in individual cells, but the trend remains the same – the averaged amount of Dps (normalized per cell) is increased upon prolonged starvation, despite the decrese in the number of survived cells (Table 1).

**Table 1.** Viable cell number decrease during starvation. The number of vegetative and dormant cells *E. coli* Top10/pBAD-DPS per 1 ml grown on LB medium with and without Dps overexpression.

| 6 hrs |  | 24 hrs |  | 48 hrs |  | 2 months |  | 7 months |  |
| --- | --- | --- | --- | --- | --- | --- | --- | --- | --- |
| $1.9 \cdot 10^9$ | $4.0 \cdot 10^9$ | $1.4 \cdot 10^9$ | $3.6 \cdot 10^9$ | $7.4 \cdot 10^8$ | $1.8 \cdot 10^9$ | $5.3 \cdot 10^7$ | $2.5 \cdot 10^7$ | $2.9 \cdot 10^7$ | $6.9 \cdot 10^6$ |
| + | - | + | - | + | - | + | - | + | - |
| induction | induction | induction | induction | induction | induction | induction | induction | Induction | induction |

### Three types of condensed DNA-Dps structures found in dormant *E. coli* cells

Using transmission EM and electron two-axis tomography, we visualized and analyzed three distinct types of DNA-Dps condensation in dormant *E. coli* cells, starving up to 7 months: *nanocrystalline, liquid crystalline*, and, the most intriguing in our opinion, the *folded nucleosome-like* type. The first two types were extensively described previously [3,4,6,15,19,21], but the third one, to our knowledge, is novel, and has been observed here for the first time in the cells, starving for up to 7 months. The *nanocrystalline* are highly ordered intracellular structures of various sizes and shape, that often form 3D arrays, built from layers of DNA, stabilized by the Dps (Fig. 2A) [3,6,34]. In *liquid crystalline*, less condensed DNA is found in the central part of the cell, surrounded by a condensed cytoplasm (Fig. 2B) [3]. The *folded nucleosome-like* condensate is an abundance of spherical particles with the diameter of 30-50 nm in the cytoplasm (Fig. 2C). Sometimes, several types of the above-mentioned structures were co-detected (data not shown).

**Figure 2.**
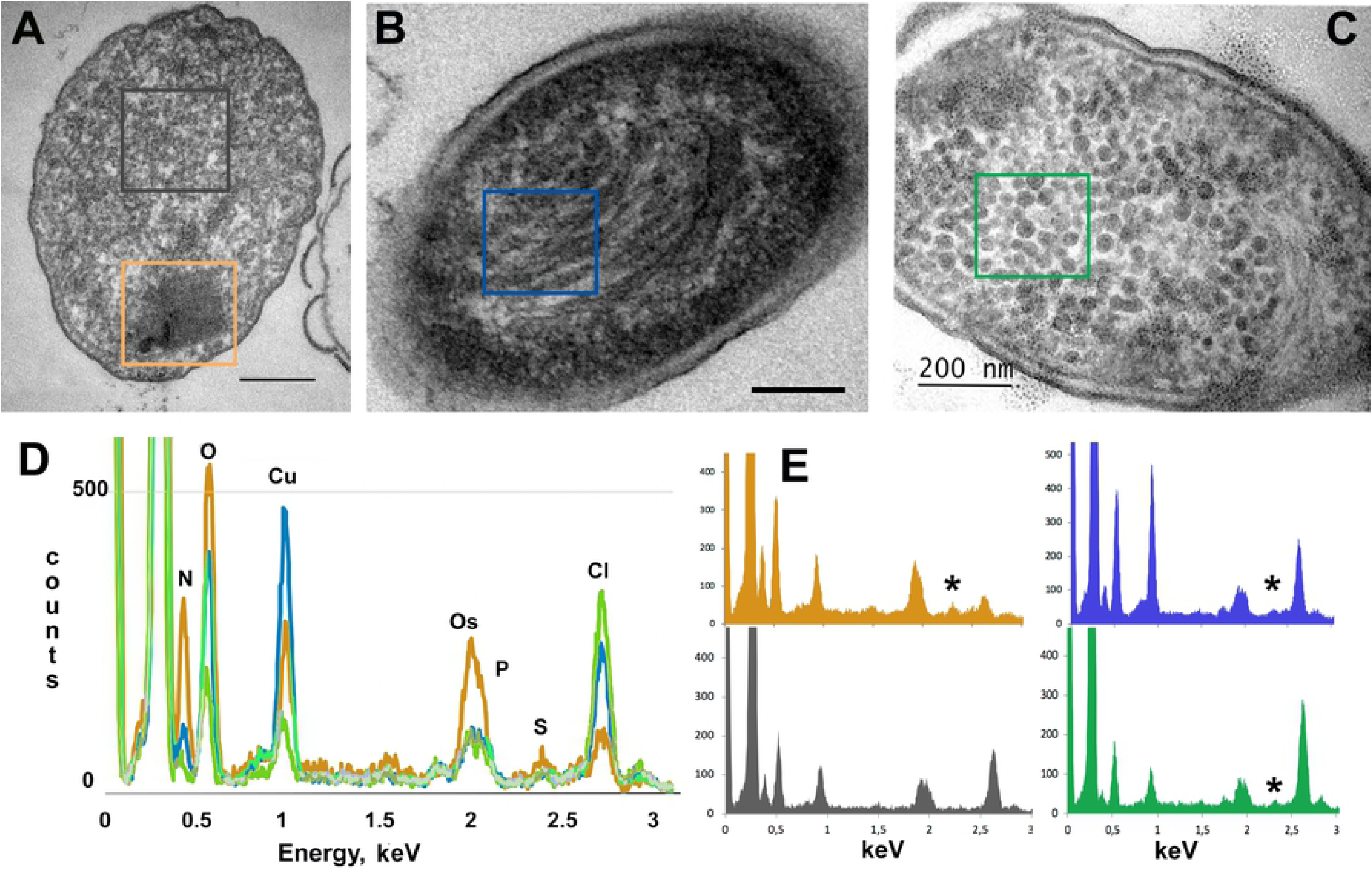
Three types of condensed DNA-Dps structures found in dormant *E. coli* cells, starving for 7 months. (A) *nanocrystalline*; (B) *liquid crystalline*; (C) *folded nucleosome-like* type. Bar – 200 nm. Colored frames mark the specific areas, where the EDX spectra were obtained; (D) Superimposed EDX spectra from the selected areas, marked in (A-D), normalized to the C peak. Line colors reflect the corresponded frame colors in (A-D). (E) EDX spectra from the selected areas marked in (A-D), colored by the specific area. Asterisks mark positions of S peak.

We confirmed the presence of both Dps and DNA in all ordered structures by EDX analysis (Fig 2D, E). This method allows to detect and map various elements on thin slices [35]. When it was possible to detect the K_a_ peak (2.307 keV) of S, we suggested that it reflected the existence of Dps (each dodecamer of Dps contains 48 Methionines), while the K_a_ peak (2.013 keV) of P corresponds to the DNA. The areas that were subjected to EDX analysis are marked on Fig. 2A-C by colored outlines. All spectra were normalized to the C peak and superimposed to each other in Fig. 2D. The pronounced Cu signal comes from the copper grids. In the reference area surrounding the ordered structure, neither S nor P have been identified (Fig. 2A, dark gray rectangle; Fig. 2D). It should be noted that the K_a_ peak (2.013 keV) of P is very close to the M-line (1.914 keV) of Os, that is used for fixing the cellular membranes.

Comparison of EDX spectra obtained for the three different ordered structures (Fig. 2D) revealed increased levels of both P and S in *nanocrystalline*, comparing to other two formations, suggesting that in the *nanocrystal* the majority of Dps protein is tightly bound to nucleoid DNA, forming a compact arrangement, which is in accordance with previous studies [21]. In *liquid crystalline* and *folded nucleosome-like* formations, somewhat smaller S peaks were also clearly detected (Fig. 2D, E, asterisks), while P peaks have the same highs as the Os peaks have. This may suggest that only a part of nucleoid DNA and some of cellular Dps population are incorporated in these more freely ordered structures.

### Morphology of condensed DNA-Dps structures in dormant *E. coli* cells

In cells with overproduction of the Dps, most of the detected structures are ***nanocrystalline***. To prove this, we generated Fourier transforms from these structures (Fig. 3A, insert; Fig. 3C). To estimate the interlayer distance, double-axis tomography (Fig. 3C) was employed. This allowed us to obtain an improved Fourier transform and, by inverse Fourier transform, an image of a DNA-Dps nanocrystal filtered from noise (Fig. 3D). The distance between the centers of Dps molecules in neighboring layers was 9 nm, which is comparable with results obtained *in vitro* [34], in cryo-EM [22], and in synchrotron radiation scattering [2].

**Figure 3.**
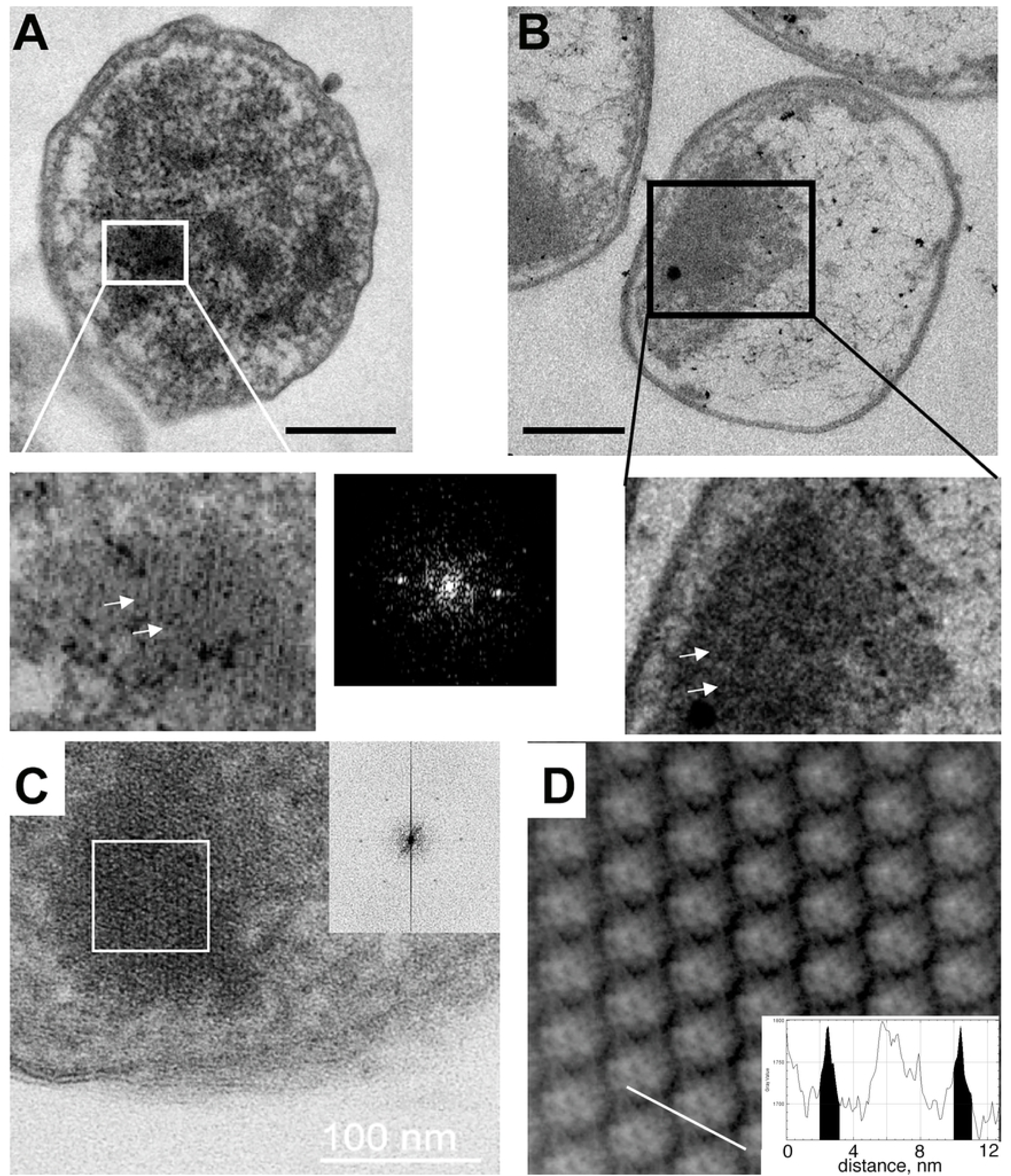
Morphology of DNA-Dps nanocrystals in dormant *E. coli* cells. (A) *E. coli* strain BL21-Gold(DE3)/pET-DPS growing on M9 media with induction of Dps overexpression, induced in the linear growth phase, age 7 months; insert – *nanocrystalline* assembly, right – FFT from the selected in (A) area; (B) *E. coli* strain Top10/pBAD-DPS growing on LB media without inducing Dps overexpression, age 7 months. Insert – amorphous DNA-Dps assembly; arrows are pointing to the individual DNA-Dps layers. Bar size – 250 nm; (C) center slice through the tomogram of the *E. coli* cell (semi-thick section) with the *nanocrystalline* structure inside; insert – Fourier transform from the white-bordered area; (D) filtered DNA-Dps co-crystal. Insert – the intensity profile along the white line on the main image. Highlighted in black are densities, corresponded to the inter-layer DNA strands.

On the other hand, in cells lacking Dps overexpression (Fig. 3B, insert), the condensed DNA-Dps structures were less ordered, which precluded us to obtain the Fourier transform from these samples. The interlayer distance in such structures dispersed from 7 to 10 nm.

***Liquid crystalline*** structure is the second common type of a DNA condensed structure that we found in dormant *E. coli* cells. The structures of this kind has been demonstrated previously in starved bacteria lacking the *dps* gene [3], in viruses, and spores of various origin [25]. In some bacterial viruses, the double-stranded DNA is stored inside the capsid in the form of a spool [36,37], which can have different types of coiling leading to different types of liquid-crystalline packaging [37-39]. This packaging can change from hexagonal, to cholesteric, to isotropic at different stages of the viral life cycle.

Each of the *E. coli* populations studied by us possessed the *dps* gene. In some cells, the DNA has the form of a cholesteric *liquid crystalline* order (Fig. 4 A, D). The DNA packaging in this condensed phase reduces the accessibility of DNA molecules to various damaging factors, including irradiation, oxidizing agents, and nucleases [3]. The cell, displayed on Fig. 4(B), possess an isotropic DNA condensation type [37-39], which is also characteristic for bacterial spores [25]. The DNA-Dps in the cell on Fig. 4C has a nearly cholesteric order. A small amount of S (Fig. 2D, E, asterisks) reflects the smaller Dps-to-DNA ratio (Fig 4E) in the *liquid crystalline* formations.

**Figure 4.**
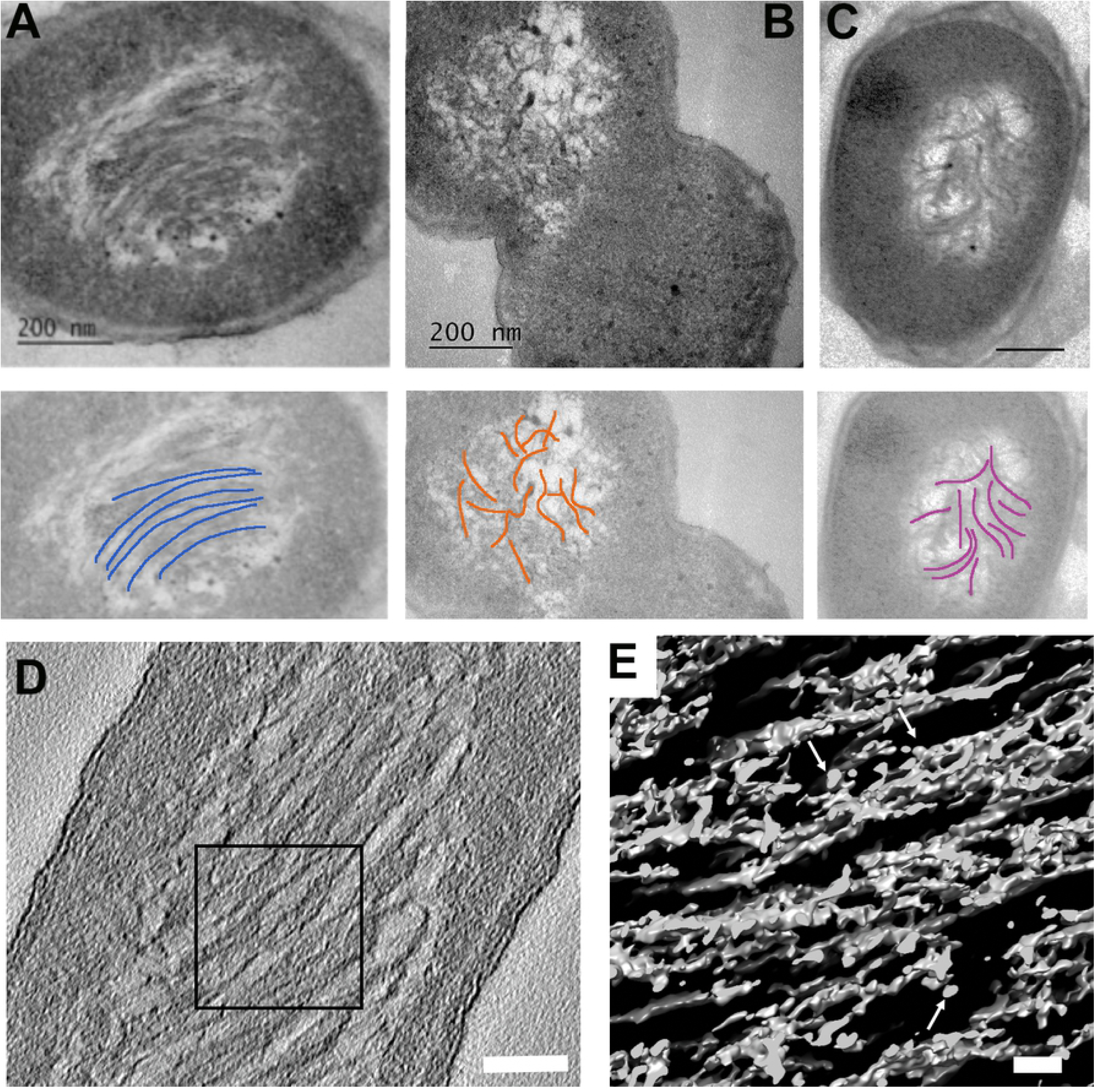
Morphology of DNA-Dps *liquid crystalline* assemblies in dormant *E. coli* cells. (A) *E.coli* strain Top10/pBAD-DPS growing on M9 media, with induced Dps production in the linear growth phase, age 7 months; (B, C) *E. coli* strain BL21-Gold(DE3)/pET-DPS, same conditions as in (A); below, each micrograph – corresponded schematic of DNA strand distribution (blue – cholesteric order, orange – isotropic order, purple – nearly cholesteric order). (D) Central section through the tomogram of an *E. coli* cell with a cholesteric *liquid crystalline* structure. Bar –100 nm. (E) 3D representation of the *liquid crystal*, marked on (D) with a black frame. White arrows are pointing to the Dps. Bar – 30 nm.

Of particular interest is the third type of ordered structure, found in dormant *E. coli* cells: the ***folded nucleosome-like***. In all of the studied populations, both with and without overproduction of Dps, cytoplasm of 5% to 25% cells was overfilled with many round structures (Fig. 5), with an average diameter of 30 nm.

**Figure 5.**
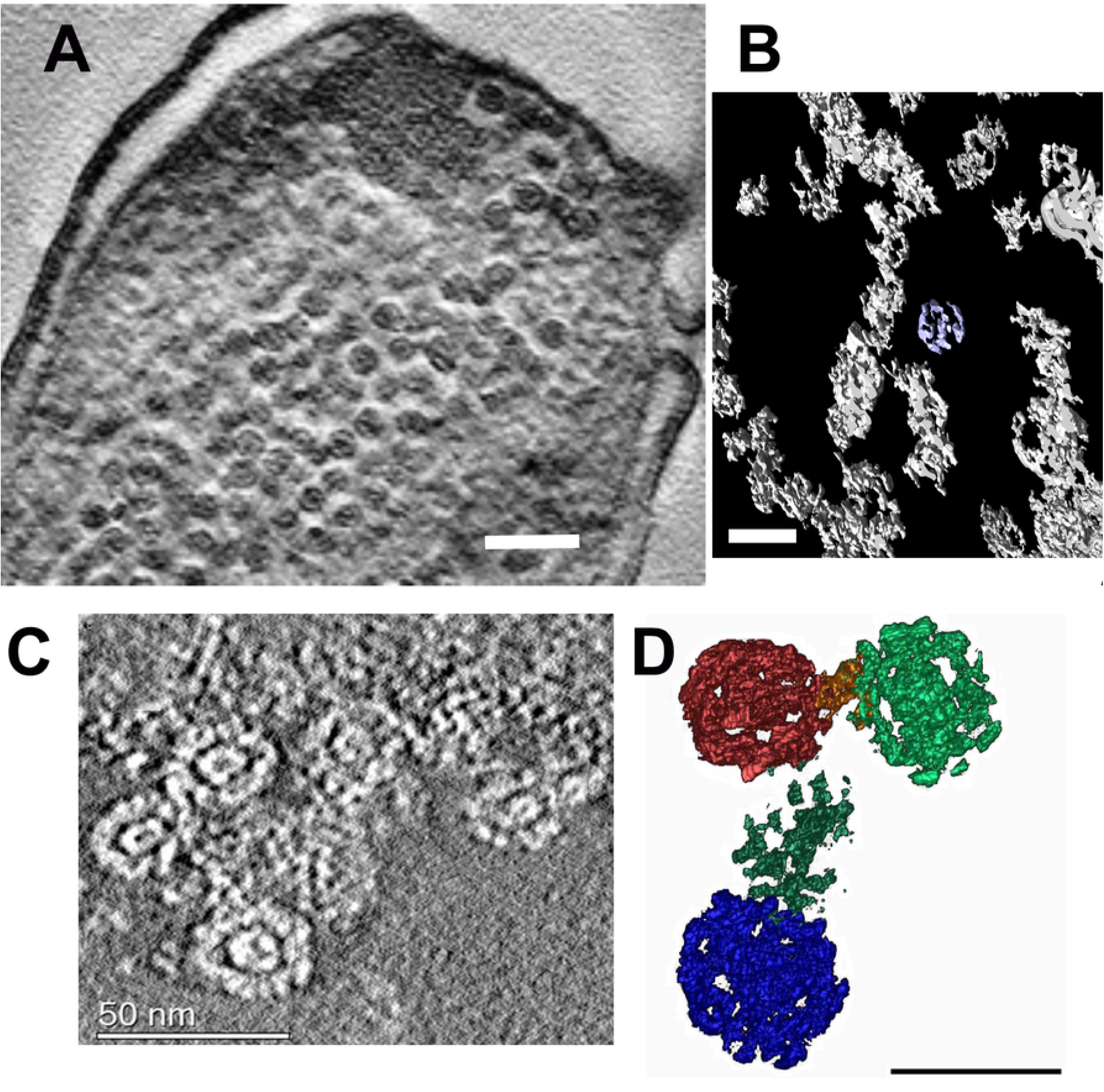
Morphology of the DNA-Dps folded *nucleosome-like* structure in dormant *E.coli* cells. (A) Tomogram of an *E. coli* cell, strain BL21-Gold(DE3)/pET-DPS growing on M9 media, with induced Dps production in the linear growth phase, age 7 months. Bar – 100 nm; (B) 3D subtomogram reconstruction of corrsponded Dps spherical associates. Bar – 50 nm; (C) Tomogram of *E. coli* cell, strain Top10/pBAD-DPS, growing on M9 media, without induction of Dps production, age 7 months. Bar size – 50 nm; (D) 3D subtomogram reconstruction of corresponded Dps spherical associates. Bar size – 30 nm.

Such structures were more often presented in cells that grew on a synthetic media (Table 2). The tomographic study (Fig. 5) clearly demonstrated that these structures are not toroids, described previously [21,40,41], but represent nearly spherical formations. Remembering that the bacterial nucleoid represents an intermediate engineering solution between the protein-free DNA packaging in viruses and protein-determined DNA packaging in eukaryotes [25], we called this type of structure the ‘*folded nucleosome-like’*.

**Table 2.**
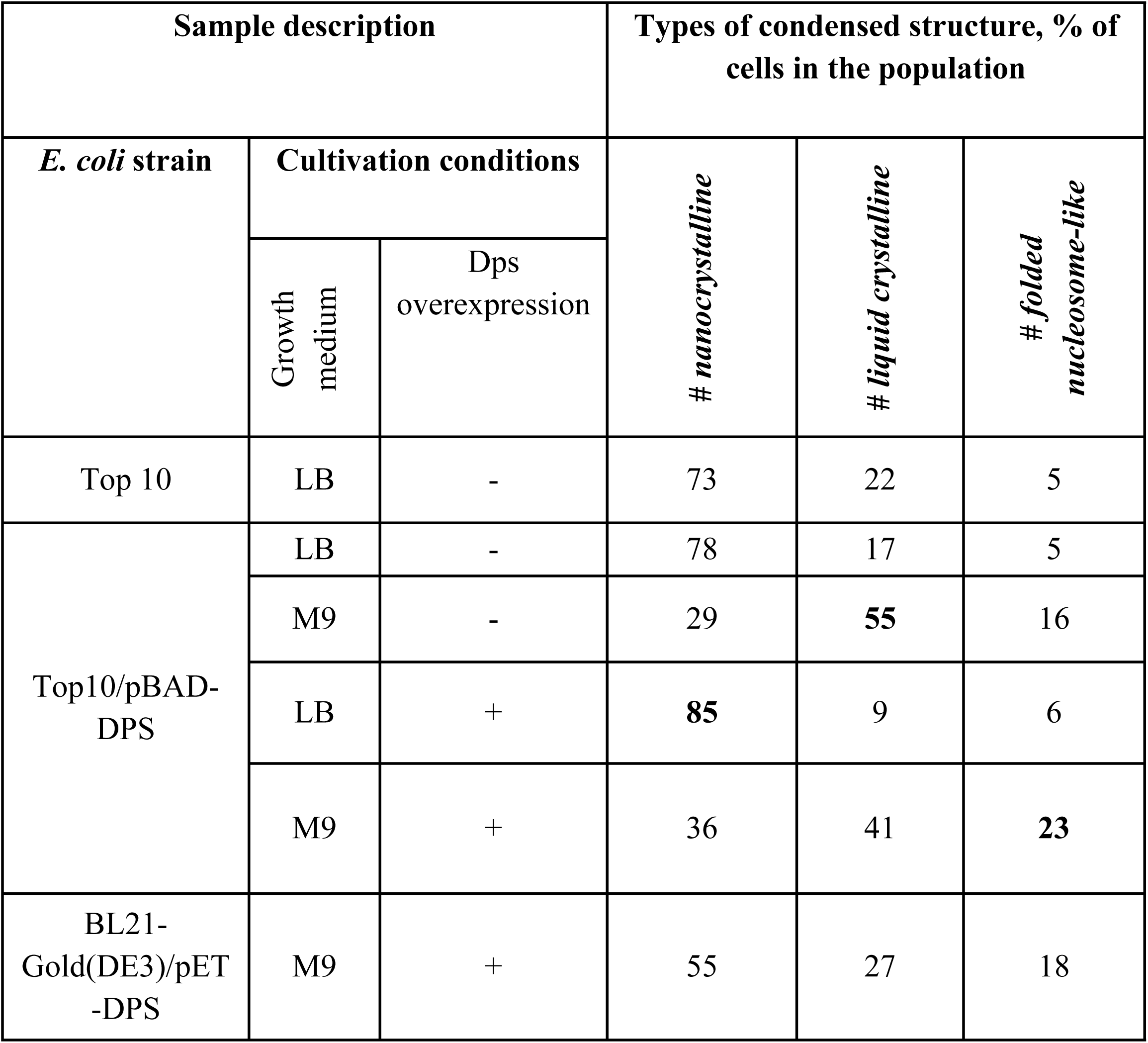
Variety of condensed DNA-Dps types in studied *E. coli* dormant cells. **Table 2** reflects a trend (as a percentage of the total number of cells) for the formation of a specific condensed DNA structure type for the cells of the certain strain and growth conditions after 7 months of cultivation.

| Sample description |  |  | Types of condensed structure, % of cells in the population |  |  |
| --- | --- | --- | --- | --- | --- |
| *E. coli* strain | Cultivation conditions |  | # *nanocrystalline* | # *liquid crystalline* | # *folded nucleosome-like* |
|  | Growth medium | Dps overexpression |  |  |  |
| Top 10 | LB | - | 73 | 22 | 5 |
| Top10/pBAD-DPS | LB | - | 78 | 17 | 5 |
|  | M9 | - | 29 | **55** | 16 |
|  | LB | + | **85** | 9 | 6 |
|  | M9 | + | 36 | 41 | **23** |
| BL21-Gold(DE3)/pET-DPS | M9 | + | 55 | 27 | 18 |

Element analysis demonstrated that the spherical aggregates, indeed, contained S and P peaks (Fig. 2D, E), indicating the presence of DNA-Dps associates.

We suggested that, in bacterial cells, spherical formations of Dps molecules (see Fig. 5B, D) either may act similarly to histones, on which the DNA is twisted (histone-like behavior) (Fig. 6B), or, which is more probable, the DNA arranges the Dps beads through which its string passes, forming ‘beads on the string’ (Fig. 6A). To counteract external stress factors, these formations should be placed quite tightly on the nucleoid DNA. In addition, like in the case of protein-determined DNA packaging in eukaryotic cells where the nucleosomes are folded to form fiber loops, which then form the chromatid of a chromosome, beads on the string (or histones) may fold into a compact structure like a globule (Fig. 6C). Possible schematic representation of the *folded nucleosome-like* structure generation suggested here may be seen on Fig 6. External adherent Dps molecules can additionally protect the DNA (Fig. 6 E).

**Figure 6.**
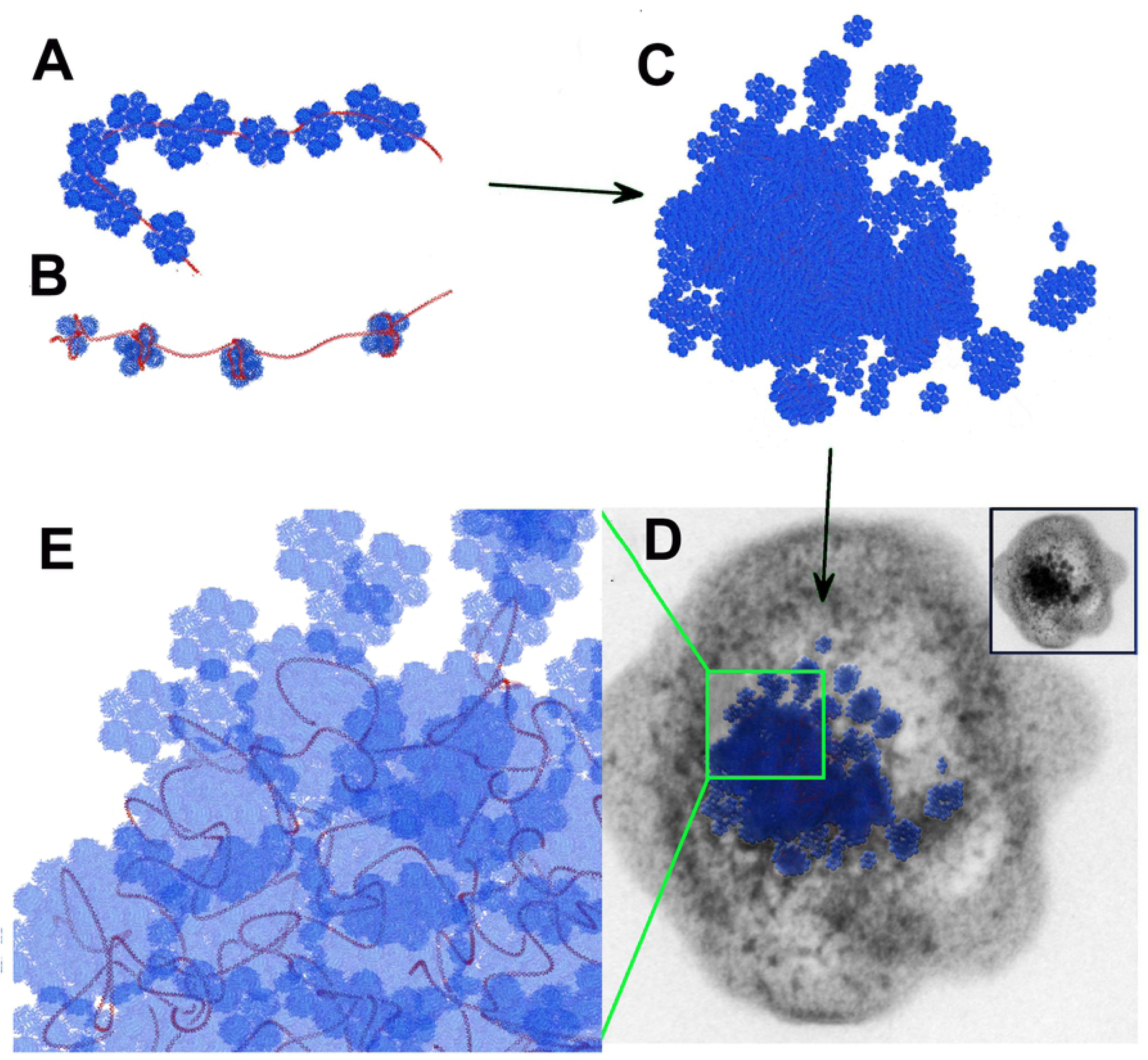
The suggested model of the *folded nucleosome-like* structure arrangement. (A) A possible folding –’beads on the string’ where the string is the DNA and the beads consist of Dps molecules (blue) as in Fig. 5D; (B) Other possible folding - spherical aggregates of Dps act similarly to histones, on which the DNA is twisted (histone-like); (C) For better protection of DNA from the external environment, the chain (beads on the string or histone - like) is folded into a globule-like structure. (D) A full image of the nucleosome-like globule superimposed with the bacterial cell. The insert – an original micrograph of the same cell. Part of the globule-like structure with adherent Dps molecules (red) is shown in (E).

### Dormant *E. coli* cells morphology indicates heterogeneity

The presence of specific ordered assemblies in dormant *E. coli* cells varied significantly depending on the strain and cultivation conditions (Table 2). That is, in the population of dormant cells from strains Top10 (wild type) and Top10/pBAD-DPS without induction of Dps overproduction (similar to the wild type), cultivated on LB medium, the cells with the *nanocrystalline* structure were predominant (more than 70%). Induction of Dps overproduction in the latter strain led to the increase of the cell number up to ∼85%. This is in accordance with earlier observations of bacterial cells starving for up to 48 hrs [2–4,15,24]. *Nanocrystalline* was the most frequently ordered structure in these cells, which consequently made them stable against the nucleases and oxidants [3,6]. Here, we observed this type of ordered structure, for the first time, in dormant cells, starving for as long as 7 months.

When the Top10/pBAD-DPS cells, without overexpression of Dps, were grown on a synthetic M9 medium (less nutrients), the number of cells with *nanocrystalline* structures decreased to 40%, but more cells with *liquid crystalline* structures appeared. In the same strain, with the overexpression of Dps, growing on synthetic M9 medium (less nutrients), a maximum number of *folded nucleosome-like* structures were detected (23%). This suggests that the presence of nutrients plays a key role in DNA compactization. In the absence of nutrients, even with high concentrations of Dps (Fig. 1), cells cannot form quality *nanocrystalline* structures.

Indeed, the sizes and the number of *nanocrystalline* structures in dormant cells vary, depending on the *E. coli* strain and level of Dps production. For example, in the strain BL21-Gold (DE3)/pET-DPS, with Dps induction, yet growing on M9 media, the number of nanocrystals usually varied from 5 to 10 per cell; with sizes of approximately 40-80 nm (Fig. 3A). In the Top10/pBAD-DPS cells, without Dps induction, grown on an LB medium and aged 7 months, one large crystal (300-400 nm size) often occupied a large part of one cell (Fig. 3B). The number of cells bearing *liquid crystalline* structures in populations of dormant *E. coli* cells ranged from ∼10% up to ∼55% (Table 2), with the majority being observed in those cells growing on M9 synthetic medium.

## Conclusions

Here, we found out that there is no single manner of DNA condensation into the heterogeneous population of dormant *E. coli* cells, starved up to 7 months. We have observed three types of ordered structure morphology, namely: *nanocrystalline, liquid crystalline*, and *folded nucleosome-like*. The fact that cells bearing the above-mentioned ordered forms were found in aged populations proves that they are not transitional intermediate forms, but that they are programmed into the development cycle for the purpose of implementing different survival strategies. The DNA does not form a unique compact structure upon condensation. The complex structure of a condensed high polymer like the DNA with a Dps protein may consist of various regions, each with a characteristic degree of the internal order [42] ranging continuously from some of it close to the ideal crystals, to a completely amorphous state. Earlier we studied dormant *E.coli* cells using synchrotron radiation diffraction [2,4]. Broad diffraction peaks found there were indicative of the imperfection of DNA-Dps co-crystals.

The heterogeneity of cells allows to respond flexibly to environmental changes and to survive in stressful situations. The main reason why we observed several types of DNA condensation in dormant *E. coli* cells – is their heterogeneity. It increases the ability of the whole population to survive under diverse stress conditions. The results, observed here, shed a new light, both on the phenomenon of DNA condensation in dormant prokaryotic cells and on the general problem of developing a response to the stress.

## Author Contribution

N.L., A.M., Y.D., G.I.E. - preparation of the samples; A.M., N.L., Y.D., O.S.S. - designing experiments, collecting, analyzing and interpreting TEM, electron tomography, and analytical EM data; K.T., V.K. - performing all calculations and statistical analysis; N.L., O.S.S., Y.F.K. - conceptualization and writing the draft of the manuscript. All authors have read and revised the draft.

## Acknowledgements

Authors would like to thank Prof. Vassili N. Lasarev for providing all *E. coli* strains, Evgeniy Kulikov and Alla Golomidova for help with gel electrophoresis and Lisa Trifonova for proofreading the manuscript. Transmission electron microscopy, analytical electron microscopy and electron tomography experiments were supported by RSF (19-74-30003). K.T., V.K., Y.F.K., N.L. and G.I.E. acknowledge the support from The Ministry of Science and Higher Education of the Russian Federation (state assignments #0082-2019-0015 and #AAAA-A19-119021490112-1). The computations were carried out on MVS-10P at Joint Supercomputer Center of the Russian Academy of Sciences (JSCC RAS). Analytical electron microscopy and dual-axis tomography were performed at the User Facility Center ‘Electron microscopy in life sciences’ of Moscow State University (Unique equipment setup ‘3D-EMC’ of Moscow State University, supported by The Ministry of Science and Higher Education of the Russian Federation, identifier RFMEFI61919×0014).

